# Polarized Dishevelled dissolution and condensation drives embryonic axis specification in oocytes

**DOI:** 10.1101/2021.05.17.444558

**Authors:** S. Zachary Swartz, Tzer Han Tan, Margherita Perillo, Nikta Fakhri, Gary M. Wessel, Athula H. Wikramanayake, Iain M. Cheeseman

**Affiliations:** Whitehead Institute for Biomedical Research, Cambridge, MA, United States; Department of Physics, Massachusetts Institute of Technology, Cambridge, MA, United States; MCB Department, Brown University, Providence, RI, United States; Department of Biology, University of Miami, Coral Gables, FL, United States; Embryology Course: Concepts and Techniques in Modern Developmental Biology, Marine Biological Laboratory, Woods Hole, MA, United States

## Abstract

The organismal body axes that are formed during embryogenesis are intimately linked to intrinsic asymmetries established at the cellular scale in oocytes [1]. Here, we report an axis-defining event in meiotic oocytes of the sea star *Patiria miniata*. Dishevelled is a cytoplasmic Wnt pathway effector required for axis development in diverse species [2-4], but the mechanisms governing its function and distribution remain poorly defined. Using time-lapse imaging, we find that Dishevelled localizes uniformly to puncta throughout the cell cortex in Prophase I-arrested oocytes, but becomes enriched at the vegetal pole following meiotic resumption through a dissolution-condensation mechanism. This process is driven by an initial disassembly phase of Dvl puncta, followed by selective reformation of Dvl assemblies at the vegetal pole. Rather than being driven by Wnt signaling, this localization behavior is coupled to meiotic cell cycle progression and influenced by Lamp1+ endosome association and Frizzled receptors pre-localized within the oocyte cortex. Our results reveal a cell cycle-linked mechanism by which maternal cellular polarity is transduced to the embryo through spatially-regulated Dishevelled dynamics.

## RESULTS AND DISCUSSION

### Dishevelled is required for specification of the primary body axis in sea star embryos

Developmental determinants of the primary body axis are maternally established within the oocytes of diverse animal species [5]. The Wnt/beta catenin pathway is an important determinant of the organismal body axis, and provides a potential link between oocyte asymmetries and the embryonic body axes [6-8]. However, the mechanisms by which the Wnt pathway becomes asymmetrically activated within the embryo to direct axis development are poorly defined. Dishevelled (**Dvl**) is an essential cytoplasmic transducer of Wnt signaling [9-12]. Thus, we sought to test whether Dvl could serve as a developmental determinant of the primary body axis.

To define the localization and function of Dvl in body axis specification, we used the sea star *Patiria miniata*, which provides a tractable system for the study of meiosis and embryogenesis [13]. In the sea star and other echinoderms, the cytologically-defined Animal-Vegetal axis of the oocyte corresponds to the Anterior-Posterior axis of the embryo (**Fig. 1A**) [5]. We found that Dvl depletion using morpholino injection disrupted the onset of gastrulation and the expression of known Wnt-beta catenin target genes in the posterior embryo (**Fig. 1B,C**) [14], consistent with an essential role for Dvl in primary axis specification. This phenotype was rescued by expression of a knockdown-insensitive Dvl-GFP fusion construct (**Fig. 1B,C**). Dvl is therefore critical for the specification of the primary body axis in sea star embryos.

**Figure 1.**
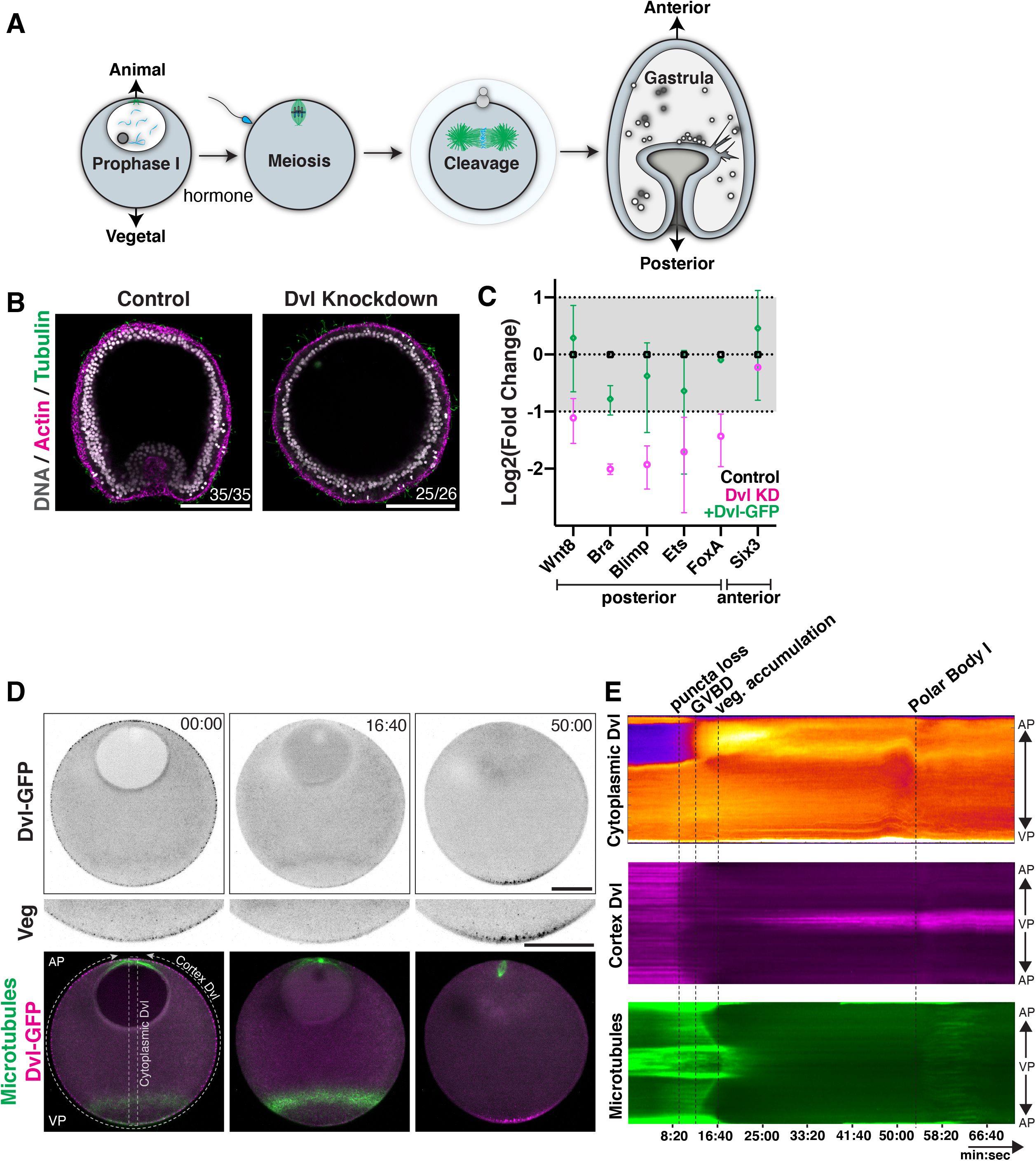
Dishevelled is required for posterior identity and undergoes relocalization in meiosis. (A) Schematic of sea star meiosis and development. The Animal-Vegetal axis in oocytes, defined by the site of the meiotic divisions, correlates with the Anterior-Posterior axis in the embryo. (B) Depletion of Dvl by morpholino injection disrupts primary invagination at the posterior pole (control n=35, Dvl knockdown n=26 embryos). (C) Expression of posteriorly expressed cWnt pathway genes is downregulated with Dvl knockdown as measured by qPCR. The anteriorly expressed factor Six3 is unaffected (mean and standard deviation of 2 biological replicates each performed in technical triplicate). (D) Live oocyte expressing Dvl-GFP and EMTB-mCherry. In Prophase I arrest (t=00:00), Dvl localizes to uniformly distributed puncta at the cortex. In addition to the centrosomal microtubule array at the animal pole, EMTB-mCherry revealed a band of microtubule density at the vegetal pole. Following hormonal stimulation, Dvl puncta dissolve (t=16:40), and then reform at the vegetal pole (t=50:00). Scale bars = 50μ. (E) Kymograph depiction of the oocyte in (D), defined by the white dashed lines in the lower left image. A transverse kymograph of cytoplasmic Dvl, along with circumferential kymographs for cortical Dvl and microtubules are provided. Regions of interest to which the kymographs correspond are outlined in (D).

### Dishevelled localization becomes polarized in meiosis

We next sought to define Dvl localization and the mechanisms that govern its behavior. Given the essential role of Dvl in axis determination, we tested the spatial localization of Dvl during the oocyte-to-embryo transition by live imaging of Dvl-GFP relative to the microtubule marker mCherry-EMTB (**Fig. 1D, movie S1**). In Prophase I-arrested oocytes, Dvl-GFP localized to puncta uniformly throughout the cell cortex, as well as diffusely to the cytoplasm (**Fig. 1D,E**). However, following hormonal stimulation, just prior to germinal vesicle breakdown (**GVBD**), these Dvl-GFP puncta were lost (**Fig. 1D,E**, 16:40). Subsequently, Dvl-GFP assemblies re-appeared in the vegetal subcortical cytoplasm as the first meiotic spindle formed. These structures increased in prevalence as the first meiotic division completed (**movie S1**). Following the completion of meiosis, the first mitotic spindle is oriented perpendicularly with the cleavage plane parallel to the animal-vegetal axis (see also [15]). Due to the asymmetric localization of Dvl to the vegetal pole, this cleavage pattern results in apportioning the vegetal Dvl domain equally to both blastomeres where it remains asymmetrically localized (**Fig. S1C**).

To confirm the behavior of the ectopically-expressed Dvl-GFP fusion, we additionally visualized endogenous Dvl mRNA and protein distribution. Using fluorescent in situ hybridization, we found that Dvl mRNA is uniformly present throughout the oocyte and embryo (**Fig. S1A,B**). Based on immunofluorescence using antibodies generated against *P. miniata* Dvl, endogenous Dvl protein localized throughout Prophase I-arrested oocytes, but formed assemblies that are enriched at the vegetal pole in meiosis (**Fig. S1C**), consistent with our observations of Dvl-GFP. In summary, Dvl localization becomes polarized in a process temporally coupled to meiotic cell cycle progression, which could serve as a critical event in the specification of the primary body axis in embryogenesis.

### Polarized dissolution and condensation coupled to meiotic progression drives Dvl localization

We next sought to define the mechanisms that underlie Dvl re-localization to the vegetal pole, including whether this requires cytoskeletal-based cytoplasmic flow or active transport along cytoskeletal elements. We did not observe substantial changes in the bulk cytoplasmic Dvl-GFP concentration along the AV axis when viewed as kymograph over meiotic time (**Fig. 1E**), arguing against the presence of directional transport. In addition, pharmacological disruption of microtubule or actin networks did not prevent Dvl vegetal enrichment following hormonal stimulation (**Fig. S2 A,B**), suggesting that Dvl distribution occurs through alternative mechanisms.

To further define the localization behavior of Dvl, we specifically imaged the vegetal pole at higher magnification and performed image segmentation and particle tracking analyses. This analysis revealed two discrete populations of Dvl granules (**Fig. 2A, movie S2**) - one in the cytoplasm, and another in close association with the cell cortex (**movie S2, S3**). We therefore separately tracked these two populations over time to determine their differential behaviors (**movie S3**). Mean squared displacement (MSD) measurements for the two populations indicated that the vegetal cortex granules are largely immobile and achieved lower velocities, whereas the cytoplasmic granules traverse greater distances (**Fig. 2B, Fig. S2D**). However, on a population level, the movements of the cytoplasmic granules were not directionally biased, based on an analysis of the mean velocities of all particle trajectories across angles relative to the Animal-Vegetal axis (**Fig. 2C**). From these results, we conclude that polarized Dvl localization is not a result of directional transport or flow.

**Figure 2.**
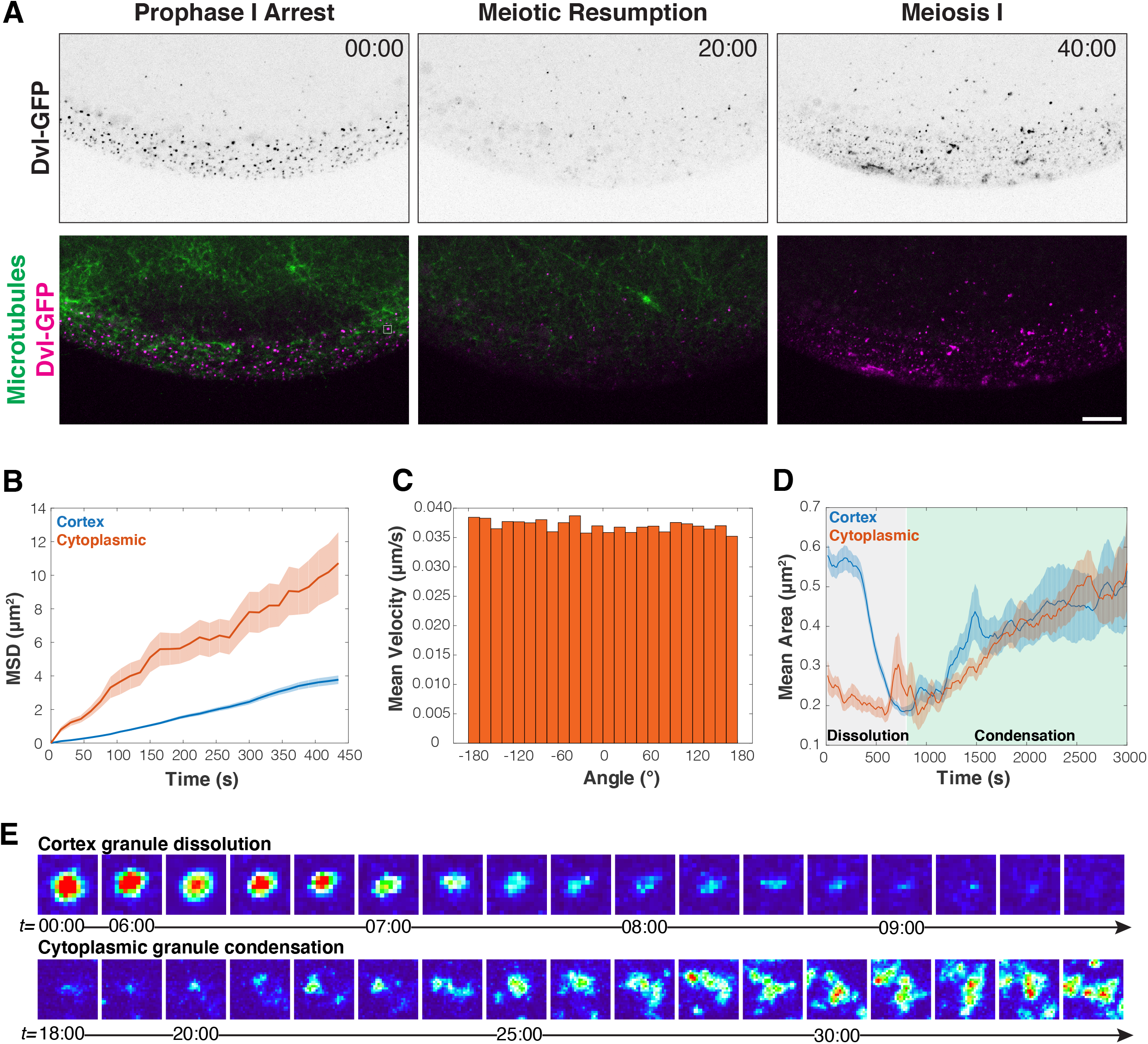
Dishevelled localization by polarized dissolution and condensation. (A) Time lapse vegetal pole images of oocyte expressing Dvl-GFP and EMTB-mCherry at high magnification (scale bar 10μ). (B) Mean squared displacement for Dvl assemblies over time, indicating cortex assemblies are less mobile. (C) Mean velocity of cytoplasmic assemblies across angles relative to the Animal-Vegetal axis reveals no directionally biased movement. (D) Area quantification of Dvl assemblies from maximally projected 3D time lapse shown in (A). The images were segmented to separately measure cortex-associated or cytoplasmic assemblies. Area measurements reveal a period of dissolution and condensation for both cytoplasmic and cortex Dvl assemblies. (B,D) Error bars represents the standard error of puncta over time, the sample size of which is provided in **Fig. S2C.** (E) Magnified views of single Dvl puncta during the dissolution or condensation period. Top row images are 1.5μ wide, and lower row images are 3.0μ wide.

We next hypothesized that biased assembly and disassembly could favor Dvl accumulation at the vegetal pole, but disfavor Dvl localization in the animal pole region of the oocyte. To test this, we measured the size of Dvl assemblies over time, using the area of maximally projected time lapse movies as a proxy. In Prophase I-arrested oocytes, the cortex-associated granules were larger and more abundant than the cytoplasmic granules. However, following hormonal stimulation, the size of cortex-associated Dvl puncta (measured by mean area) decreased rapidly, displaying behavior characteristic of dissolution (**Fig. 2D,E, Fig. S2C**). As meiosis progressed, both the cortex and cytoplasmic granules began to reform in a condensation-like process, reaching a mean area of approximately 0.5 μm^2^ (**Fig. 2D,E**). Given this strikingly dynamic disassembly and reformation in response to hormonal signaling, we conclude that Dvl localization is a result of polarized dissolution and condensation. This dissolution and condensation occurs downstream of biochemical signals associated with meiotic resumption, and is temporally coupled with meiotic progression. Together, these processes favor condensation at the vegetal pole driving asymmetric Dvl re-localization.

### Dvl granules are dynamic molecular assemblies

The rapid dissolution and reformation behavior suggests that Dvl incorporates into dynamic molecular assemblies. As a first test of these behaviors, we assessed whether Dvl granules were capable of fusion and fission. By time lapse imaging, we identified fusion events amongst the mobile cytoplasmic Dvl granules suggesting that these assemblies engage in higher-order molecular interactions (**Fig. 3A, movie S4**). To further test the nature of Dvl binding within the larger assemblies, we used fluorescent recovery after photobleaching (FRAP) to probe the cortical Dvl puncta in Prophase I-arrested oocytes, and the immobile puncta at the vegetal cortex after Dvl re-localization (**Fig. 3B,C**). We found that Dvl exchanged readily between the punctate assemblies and cytoplasmic pool with a halftime for recovery of ∼2.5 minutes for both cortical and vegetal cortex-localized granules. However, although Prophase I Dvl assemblies recovered fully (mobile fraction = 1.05+/-0.32 S.D.), the vegetal cortically-localized granules displayed lower recovery (mobile fraction = 0.66+/-0.19 S.D.) (**Fig. 3C**). These differences in FRAP recovery may reflect different binding modes for Dvl in these different contexts, such as a vegetal receptor for Dvl that is distinct from its binding mode in Prophase I-arrested oocytes. Although these data do not allow us to conclusively determine the physical nature of the phase transition state that Dvl assemblies undergo, based on the dynamic properties measured on FRAP and the ability of assemblies to fuse, we propose that Dvl assemblies are dynamic biomolecular condensates.

**Figure 3.**
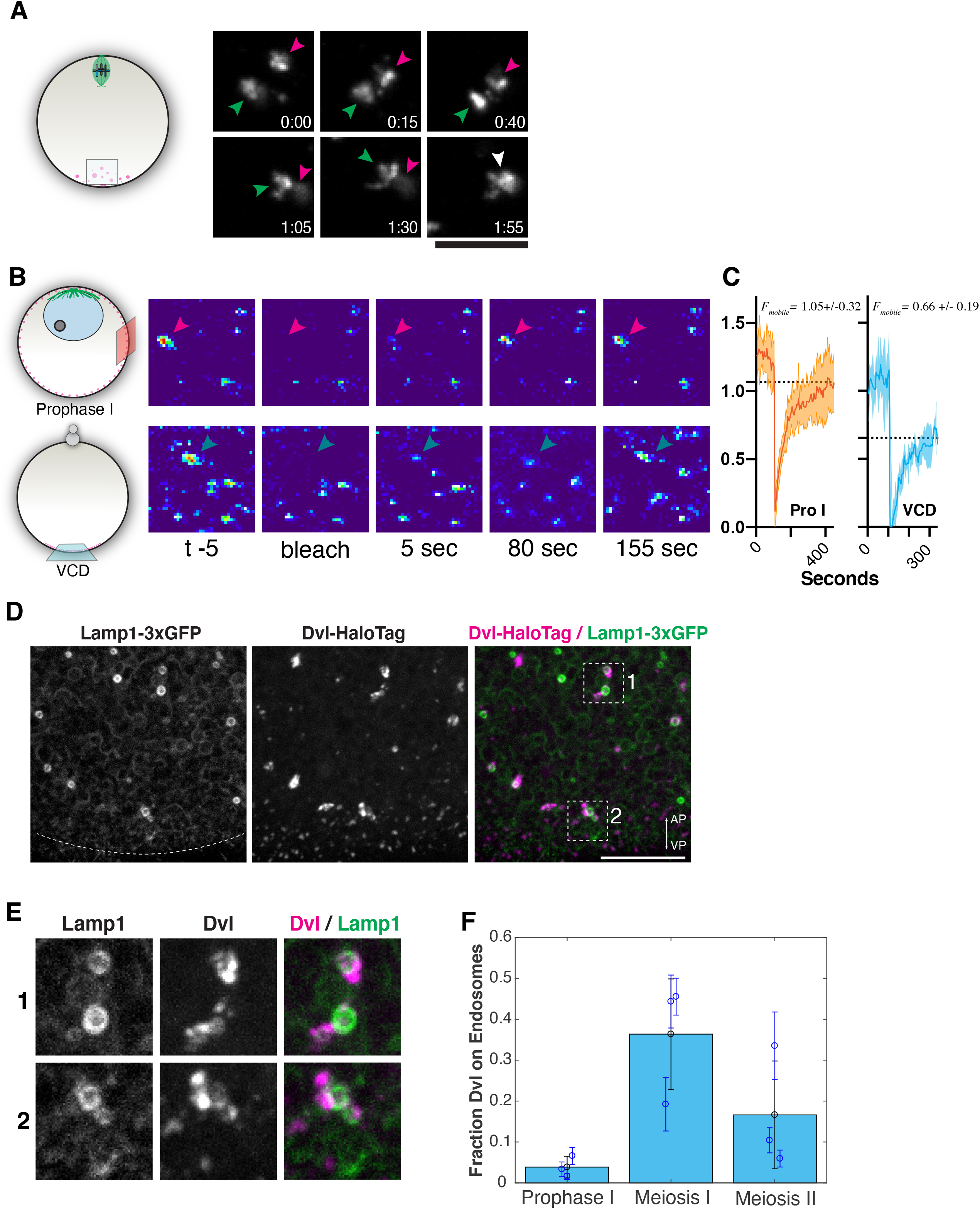
Dishevelled forms dynamic molecular assemblies and associates with cytoplasmic organelles. (A) Time lapse video stills of a cytoplasmic Dvl assembly (in area schematized to the right) undergoing fusion. Arrowheads indicate the two separate assemblies. (B) FRAP analysis of uniform cortex puncta in Prophase I, or vegetal cortical domain (VCD) puncta in meiosis. Stills from example FRAP experiments are provided. (C) FRAP quantification of Prophase I or VCD granules (Pro I n=6 oocytes, mean of 3 puncta each, VCD n=4 oocytes, mean of 3 puncta each). (D) Vegetal pole image of a live oocyte expressing Lamp1-3xGFP and Dvl-HaloTag showing cytoplasmic Dvl assemblies are associated with endosomes. (E) Zoom views of insets indicated in (D). (F) Fraction of Dvl assemblies that co-localize with Lamp1-positive endosomes in Prophase I, Meiosis I, and Meiosis II. Black circle is the mean value over N=3 oocytes while blue circle is the number from individual oocyte. Error bars represent the standard deviation over 10 minute time lapse videos.

### Dvl assemblies associate with Lamp1-positive endosomes

The molecular interactions and cellular states that trigger the self-association of Dvl into punctate structures are not well understood [16]. The sea star oocyte provides a temporally-controlled system in which granule formation is readily triggered by hormonal stimulation, leading to the formation of large Dvl assemblies within a single cell. The distinct Dvl populations display differing properties, which could reflect different binding partners or differing microenvironments associated with distinct cellular compartments. The movement behaviors and geometry of cytoplasmic Dvl assemblies suggest that they could be influenced by interactions with cellular structures (**Fig. 2C, 3C, movie S2**). To test this, we visualized endomembrane compartments, including putative lysosomes as labeled with Lamp1-3xGFP. We observed a striking association between Dvl granules and the surface of Lamp1-positive endosomes that was stably maintained as Dvl granules changed in shape and the endosomes moved (**Fig. 3D,E, movie S5**). Dvl-associated endosomes were also Rab7 positive, suggesting that Dvl associates with a subclass of late endosomes or lysosome-related organelles **(Fig. S3**). This unexpected interaction between Dvl granules and Lamp1-positive endosomes could facilitate nucleation of Dvl assemblies, or serve as anchors to retain them at the vegetal pole of the oocyte. Based on image segmentation, the relative proportion of Dvl in granules associated with endosomes is low in Prophase I and peaks in meiosis I (**Fig. 3F**), indicating regulation at the cell cycle level. We hypothesize that assembly of Dvl on the surface of these endosomes could provide a reservoir of Dvl protein that stably maintains the vegetal Dvl domain as the cortex is remodeled across the cell cycle.

### Pre-localized cues are required for vegetal Dvl assembly

The process of Dvl redistribution described above occurs in single oocytes denuded of associated somatic cells in open seawater. Because this would preclude Wnt signaling from an adjacent cell, we reasoned that Dvl must be cell-autonomously regulated within the oocyte. One possibility to explain this would be a gradient of diffusible factors that influence Dvl behavior. This model is analogous to MEX-5 regulation of P-granules in *C. elegans*, or the CyclinB-Cdk1 activity gradient directing the contraction wave in sea star oocytes [17, 18]. In this model, a repressive signal from the animal pole could prevent Dvl condensation except at the point farthest from its source. Importantly, such a gradient system would be sensitive to changes in oocyte geometry. Therefore, to test this model, we used microfabricated oval-shaped PDMS chambers to alter oocyte shape by shortening the Animal-Vegetal axis (**Fig. S4A**)[19, 20]. If proximity to the animal pole represses Dvl granule formation, this shape change would cause Dvl to condense at the corners of the oval - the points farthest from the animal pole. Instead, we found that Dvl-GFP localized to the original vegetal pole within the microtubule ring (**Fig. S4B**). This suggests that a cortical cue, rather than an activity gradient, regulates the re-condensation of Dvl at the vegetal pole.

We next tested whether the vegetal cortex and adjacent cytoplasm contain pre-established signals that enable Dvl condensation (**Fig. 4A**). Using microdissection, we removed the vegetal portion of the oocyte during prophase I arrest, prior to Dvl polarization (**Fig. 4A**). As a control, we removed an equivalent amount of the lateral side of the oocyte cortex and cytoplasm. In side-cut control oocytes, Dvl localized to the vegetal pole normally following hormonal stimulation. However, in vegetally-cut oocytes, while Dvl-GFP could localize to uniform cortex puncta in Prophase I, it failed to reform at the vegetal pole during meiosis (**Fig. 4A**). Reciprocally, isolated vegetal oocyte fragments displayed autonomous Dvl-GFP condensation in response to hormonal stimulation, suggesting the vegetal region of Prophase I oocytes is capable of directing Dvl condensation in the absence of other cellular structures or signals (**Fig. 4B**). Oocytes with the vegetal pole removed matured and fertilized normally, but produced embryos that failed to gastrulate, consistent with a primary axis defect (**Fig. S4C**, see also [21]). Thus, a pre-existing cue established in the vegetal pole prior to meiotic resumption is essential for the subsequent local Dvl condensation, which in turn is necessary for gastrulation of the posterior pole during embryogenesis.

**Figure 4.**
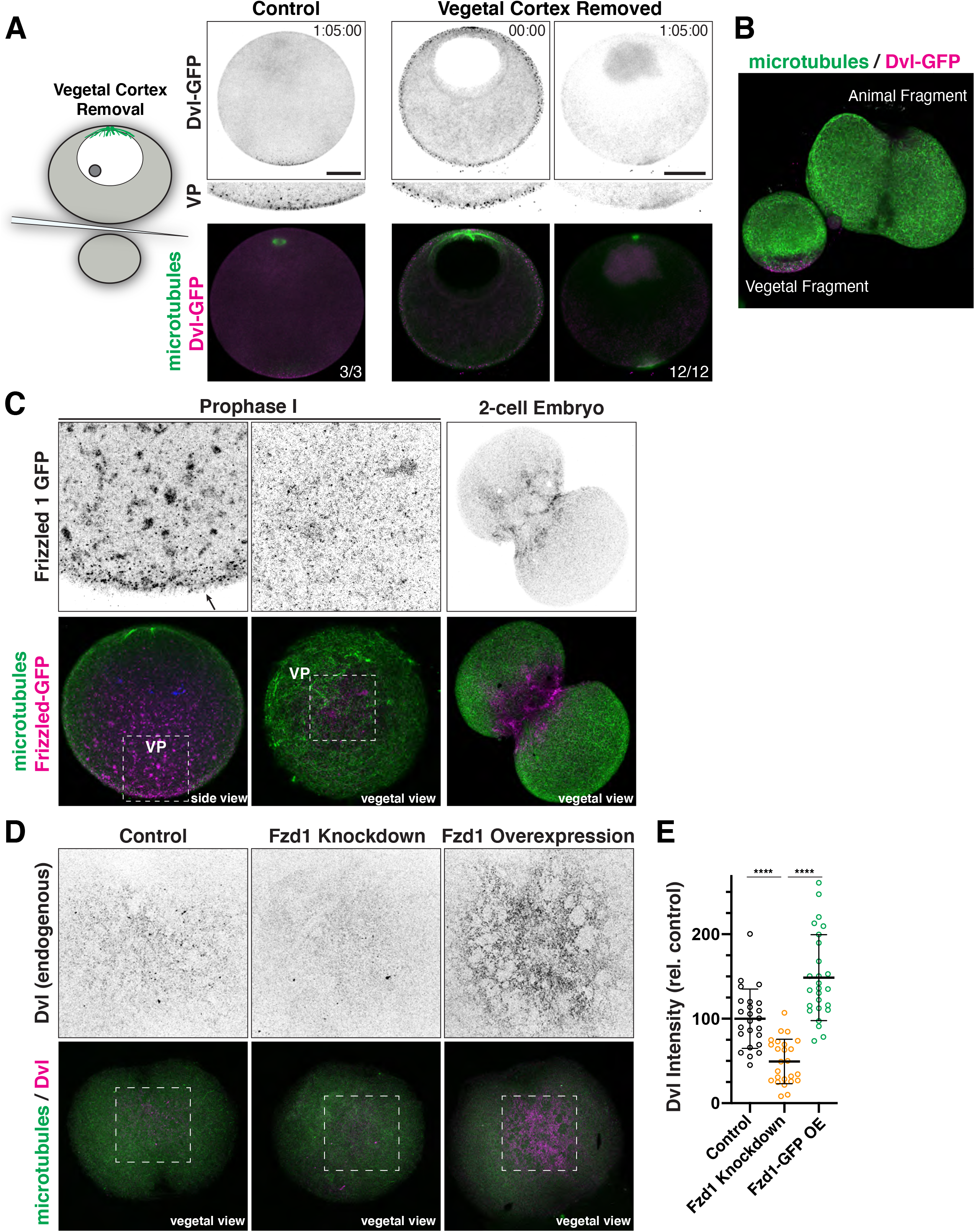
Localization mechanisms driving Dishevelled behavior. (A) Live oocytes expressing Dvl-GFP and EMTB-mCherry from which the vegetal pole was removed by microdissection as described by schematic to the right (n=12 oocytes). As a control, the side of the oocyte was removed (n=3 oocytes). (B) Animal and vegetal oocyte fragment pair following the completion of meiosis. (C) Expression of GFP-tagged Frizzled 1 in Prophase I arrested oocytes. Frizzled 1 localization is enriched to the vegetal cortex. Arrow indicates oocyte vegetal cortex. (D) Immunofluorescence images of endogenous Dvl in 2-cell embryos following control injection, Fzd1 morpholino, or morpholino combined with Fzd1-GFP overexpression. (E) Relative pixel intensity quantification of vegetal cortex Dvl signal (control n=24, Fzd1 knockdown n=25, Fzd1 overexpression n=26 oocytes, ****p < 0.0001Mann-Whitney Test).

### Maternally-expressed Frizzled 1 is required for Dvl localization

Based on the Dvl localization behavior in our oocyte microdissection experiments, we reasoned that a pre-localized membrane associated factor may serve to localize Dvl asymmetrically. Dvl can be recruited to the membrane through interactions with Frizzled transmembrane receptors, which detect Wnt signals. We therefore hypothesized that a Frizzled receptor expressed in oogenesis might be pre-established in the vegetal pole in Prophase I-arrested oocytes, prior to Dvl localization in meiosis. Analysis of a previously published transcriptome of the sea star ovary revealed the presence of Frizzled 1 (Fzd1) [22]. To test whether Fzd1 is endogenously expressed in oocytes, we assessed localization of its mRNA by FISH, and found that it is present in small primordial oocytes and large, full-grown oocytes (**Fig. S4D**). To test the subcellular localization of this putative receptor, we expressed Fzd1 as a GFP fusion in prophase I-arrested oocytes. Strikingly, Fzd1 localized to punctate structures in close association with the cell membrane specifically at the vegetal pole (**Fig. 4C**). Fzd1 also localized to cytoplasmic compartments likely corresponding to the Golgi apparatus, in a gradient that was strongest at the vegetal pole, possibly suggesting polarized trafficking of Fzd1 to the vegetal membrane.

We next tested whether Fzd1 promotes the localization of endogenous Dvl to the vegetal pole by maternal knockdown using morpholino injection. In contrast to controls, Fzd1-depleted embryos displayed significantly reduced punctate localization of Dvl to the membrane (**Fig. 4D,E**). Dvl localization to the membrane could be rescued and even increased by co-injecting knockdown-insensitive Fzd1-GFP with the anti-Fzd1 morpholino (**Fig. 4D,E**). Collectively, these results indicate that Fzd1 is localized maternally to the vegetal pole, and is essential for Dvl localization to the vegetal cortex. However, the localization of large cytoplasmic Dvl assemblies is Fzd1-independent, and may thereby provide a reservoir of Dvl protein for membrane association when additional Fzd1 receptors are incorporated into the membrane.

### A conserved paradigm for primary body axis specification in animals

A fundamental problem in development is the generation of asymmetries along body axes to pattern a multi-functionalized body plan. Comparative studies point to asymmetric activation of the Wnt pathway as a deeply conserved determinant of the primary body axis, in both cnidarians and bilaterian animals [6, 8]. Here, we identified a symmetry-determining event involving the Wnt pathway effector Dvl. In contrast to other polarization events influenced by sperm entry, such as in *C. elegans*, the sea star body axis is specified entirely by maternal control. We find that Dvl localizes to cytoplasmic and cortex-associated granules and undergoes a cycle of polarized dissolution and reassembly to distribute asymmetrically to the vegetal pole. This redistribution is essential for primary axis specification in development and is regulated cell-autonomously by cues, including the receptor Fzd1, that are prelocalized to the vegetal pole.

An important open question is how these cues are maternally established during oogenesis. In meiosis, Dvl serves as a key transducer to connect cellular asymmetries within the oocyte to body axis asymmetries at the embryonic scale. Dvl has been noted to form large “puncta” or “granules” in several animal models, as well as in mammalian cell culture based on overexpression experiments, but the physiological relevance and mechanisms by which these granules form has been debated [23, 24]. Here, we define a paradigm by which Dvl granules form at endogenous levels by dissolution and condensation, and resolve prior discrepancies regarding its endomembrane association [2, 4, 25]. An important future direction will be to determine the biophysical mechanisms underlying this dissolution-condensation process. The almost switch-like response of cortical Dvl dissolution dynamics (<5 mins, Fig. 1E, Fig. 2C) could result from a rapidly propagating signaling wave that lowers the condensation point of Dvl puncta across the cortex. In contrast, the gradual condensation process (>30 mins, Fig. 1E, Fig. 2C) suggests a nucleation-and-growth process that is seeded by endosomes and Fzd1 in a diffusion-limited manner. We hypothesize that Dvl assembly on lysosomes and related organelles enables a local reservoir of Dvl protein to support a stable posterior-specifying signaling domain through the meiotic and embryonic mitotic divisions. Given the broad diversity of organisms in which Dvl granule formation has been observed, Dvl and its capacity for spatially regulated condensation may occupy a critical node in the evolution of the Metazoan body plan.

## Acknowledgements

We thank the Marine Biological Laboratory and the students, faculty, and staff of the Embryology Course, where the pilot experiments for this work began. We thank Gene-Tools for donating the PmDvl morpholino to the course. We thank Hannah Rosenblatt for suggesting an examination of lysosomes. We thank Peter Lenart, Brad Shuster, Peter Reddien, and members of the Cheeseman laboratory for helpful discussions. We thank Veronica Hinman and her laboratory for sharing in situ probes. This research was supported by 1K99HD099315 to S.Z.S., R35GM126930 to I.M.C., and a grant to I.M.C. from the Gordon and Betty Moore Foundation. The Embryology course is supported by R25HD094666.

## STAR METHODS

### Lead Contact and Materials Availability

Requests for resources and reagents should be directed to and will be fulfilled by the Lead Contacts, Zak Swartz or Iain Cheeseman. Plasmids and antibodies are available on request, and plasmids will be additionally deposited to Addgene.

### Experimental Model and Subject Details

Sea stars *(Patiria miniata)* were wild-caught by South Coast Bio Marine (http://scbiomarine.com/) and kept in artificial seawater aquariums at 15° C. Intact ovary and testis fragments were surgically extracted from small incisions on the oral side of the animal using forceps and kept in filtered seawater containing 100 units/mL pen/strep solution at 15° C [26].

### Method Details

#### Ovary and Oocyte Culture

Ovary fragments were maintained in in artificial seawater containing 100 units/mL pen/strep solution. Intact ovary fragments were cultured this way for up to 1 week until oocytes were needed, with media changes every 2-3 days. Isolated oocytes were cultured for a maximum of 24 hours in artificial seawater with pen/strep. To induce meiotic resumption, 1-methyladenine (Acros Organics) was added to the culture at a final concentration of 10 μM. For fertilization, extracted sperm was added to the culture at a 1:1,000,000 dilution. Cytochalasin D was used at a final concentration of 40 μM, nocodazole was used at 3.3 μM. For knockdown of maternal Fzd1 (e.g. Fig. 4D,E), oocytes were injected with 500 μM morpholino and cultured for 2 days in sulfamethoxazole/trimethoprim-containing seawater (50 μg/ml sulfamethoxazole, 10 μg/ml trimethoprim).

#### Construct and antibody generation

*Patiria miniata* gene homologs were identified using genomics tools at echinobase.org [27]. Fluorescent protein fusion constructs for mRNA in vitro transcription were cloned into vectors derived from pCS2+8 using standard restriction enzyme-based methods [28]. *Dvl* protein corresponding to the DIX domain (residues 1-90) was expressed as an n-terminal GST fusion in the pGEX-6P vector in BL21 bacteria. Expression was induced with 0.1 mM IPTG and the cells were grown overnight at 18° C. GST-DIX was then purified from bacterial lysate using Glutathione-agarose (Sigma), eluted with reduced L-glutathione (Sigma). Using this immunogen, antibodies were then generated in rabbits (Covance). For affinity purification, serum was pre-depleted against GST, and then purified with GST-DIX bound to HiTrap NHS-activated columns (GE Healthcare).

#### Oocyte Microinjection and Manipulation

For expression of constructs in oocytes, plasmids were linearized with NotI to yield linear template DNA. mRNA was transcribed in vitro using mMessage mMachine SP6 and the polyadenylation kit (Life Technologies), then precipitated using lithium chloride solution. Prophase I arrested oocytes were injected horizontally in Kiehart chambers with 10-20 picoliters of mRNA solution in nuclease free water as described [26]. Dvl-GFP wildtype and mutant constructs were injected at 800 ng/μl. After microinjection, oocytes were cultured 18-24 hours to allow time for the constructs to translate before 1-methyladenine stimulation. Custom synthesized morpholinos, or the Gene Tools standard control, were injected at 1000 μM for Dvl and 500 μM for Fzd1 (Gene Tools). For microsurgery experiments (**Fig. 4A, S4C**), oocytes were cut by hand using a pulled capillary tube as a knife and a mouth pipette to hold in place. For shape mold experiments (**Fig. S4A,B**), chambers were fabricated by casting PDMS onto patterned silicon wafers as described previously [20]. The chamber shapes were designed with a height of 80 µm and surface area of around 27000 µm2, to match typical volumes of the oocytes. The patterned silicon wafer was manufactured using photolithography (Microfactory SAS, France). The silicon wafer was silanized with Trichlorosilane (Sigma 448931). PDMS was made by mixing Dow SYLGARD™ 184 Silicone Elastomer Clear solution at a 10:1 base-to-curing agent ratio. After mixing thoroughly, the elastomer was poured over the silicon master mold, degassed in a vacuum chamber and cured at 60 °C in an oven for one hour.

#### Immunofluorescence, In Situ Hybridization, and Imaging

For endogenous Dvl immunofluorescence (**Fig. S1C**), oocytes were fixed at various stages in a microtubule stabilization buffer (1% paraformaldehyde, 0.1% Triton X-100, 100 mM HEPES, pH 7.0, 50 mM EGTA, 10 mM MgSO_4_, 400 mM dextrose) for 15 minutes at room temperature [29]. The oocytes were then transferred to a solution of 80% methanol, 20% DMSO, and incubated on ice for at least 1 hour and up to overnight. Oocytes were then washed 3 times in PBS with 0.1% Triton X-100 (PBSTx), transferred to sodium citrate buffer (10 mM tri-sodium citrate dihydrate, 0.05% Tween 20, pH 6.0) and heated at 55° C for 1 hour for antigen retrieval. Oocytes were then washed 3 times with PBSTx and blocked for 15 minutes in AbDil (3% BSA, 1 X TBS, 0.1% triton X-100, 0.1% Na Azide). Primary antibodies diluted in AbDil were then applied and the oocytes were incubated overnight at room temperature. Anti-alpha tubulin (DM1α, Sigma) and Dishevelled (this study) antibodies were used at 1:5,000. DNA was stained with Hoechst. GFP fusion construct expressing oocytes were instead fixed with 2% PFA in Millonig’s Buffered Fixative (0.2 M NaH_2_PO_4_*H_2_O, 0.136 M NaCl, pH 7.0) overnight at room temperature, permeabilized by three washes with PBSTx, and blocked with AbDil as described above. GFP booster (Chromotek) was used at 1:1000 to improve the signal from GFP fusion constructs (e.g. **Fig. S2B, 4C**). The oocytes were compressed under coverslips in ProLong Gold Antifade Mountant (ThermoFisher).

Fluorescent whole mount in situ hybridization (FISH) was performed as previously described [30]. Embryos were collected and left in fixation solution (4% paraformaldehyde, 32.5% sea water, 32.5 mM MOPS pH 7, 162.5 mM NaCl) overnight at 4°C, then washed in MOPS and stored in 70% ethanol. After rehydration, embryos were hybridized with the Frzd1 probe (0.1 ng/μl) in hybridization buffer (70% formamide, 100 mM MOPS pH 7, 500 mM NaCl, 0.1%Tween 20, 1 mg/ml BSA.) at 60°C for a week. Signal was developed with the fluorophore-conjugated tyramide kit (Perkin Elmer, Cat. #:NEL752001KT; RRID:AB_2572409). After post hybridization washes in maleic acid buffer, embryos were blocked in the blocking buffer provided by the kit and left overnight with anti-digoxigenin-peroxidase antibody (Roche cat #11207733) diluted 1:2000. Signal was detected by staining for 30 min in 1:400 cy3 in amplification diluent (provided in the kit).

Fixed oocytes were imaged with a Nikon Ti2 microscope with a CoolSnap HQ2 CCD camera and a 100x 1.40 NA Olympus U-PlanApo objective (**Fig. S2B**), or a Zeiss 710 confocal **(Fig. S1C)**. Live confocal imaging was performed with a Yokogawa W1 spinning disk microscope (**Fig. 2-4, S1,3,4E**) or a Zeiss 710 (**Fig. S4A**). Images were processed with Fiji, and scaled equivalently across conditions unless otherwise specified [31].

#### RNA extraction and quantitative real-time PCR (qPCR)

RNA from 100 embryos was isolated with the RNeasy Micro kit (Qiagen, Cat#:74004). cDNA synthesis was performed using Maxima kit (Life Technologies, Cat#:K1641). qPCR was performed using ABI900 real time instrument with Maxima SYBR master mix (Life Technologies, Cat#:FERK0222) and normalized to ubiquitin transcripts. Experiments were run in two biological replicates, and each biological replicate was run on the qPCR machine with three technical replicates.

#### Quantification and Statistical Analysis

Statistical analyses were performed using Prism (GraphPad Software). Details of statistical tests and sample sizes are provided in the figure legends. Unless stated otherwise in the figure legends, images comparing the same signal across conditions are scaled equivalently. In all cases in which pixel intensity is quantified, unscaled images with appropriate background subtraction were used, as outlined below.

Kymograph depictions of Dvl localization dynamics (**Fig. 1E**) were generated in MATLAB. The space time kymograph of cytoplasmic Dvl *I*_*C*_*(y, t)* was obtained by extracting the fluorescence intensity *I*_*C*_*(y)* along the animal-vegetal (AV) axis for all time frame *t*. Here, we used *y* to denote the distance along AV axis. We first performed a Gaussian filtering step (with standard deviation of 2 pixels) to smooth the fluorescence image. We then computed the average of the intensity values 10 pixels to the left and to the right of the AV axis to obtain *I*_*C*_*(y)*. To construct the full kymograph *I*_*C*_*(y, t)*, the intensity *I*_*C*_*(y)* at each time frame *t* is aligned such that the animal and vegetal poles correspond to the same position and resampled at the appropriate distance *y*. The space time kymograph of cortical fluorescence *I*_*R*_*(s, t)* is computed by first extracting the boundary of the oocyte 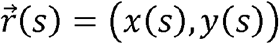 and then extracting the fluorescence intensity *I*_*R*_*(s)* along the boundary for all time frames *t*. Here, we used *s* to parameterize the arclength of the oocyte boundary. For each time frame *t*, we performed a Gaussian filtering step (with a standard deviation of 1.2 pixels) before applying a thresholding step to make a binary image. The oocyte boundary 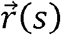 is obtained by using the *bwboundaries* function in MATLAB on the binary image. The intensity *I*_*R*_*(s)* is obtained by first identifying a local window of size 8-by-8 pixels centered at 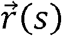 and taking the mean intensity of the pixels in the top 50 percentile intensity within the local window. To construct the full kymograph *I*_*R*_*(s, t)*, the intensity *I*_*R*_ *(s)* at each time frame *t* is aligned such that the AP corresponds to the same arclength position and resampled at the appropriate arclength *s*.

To segment the cortical and cytoplasmic Dvl puncta at a single time point (**Fig. 2C-E**), each z slice of the confocal z-stack was first demarcated into a cytoplasmic and a cortical region. By applying a thresholding to identify the boundary of the oocyte, the region around 30 pixels extending from the cell boundary into the cytoplasm is defined as the cortical region, while the rest of the oocyte is considered the cytoplasmic region. After performing this step on all z-slices, a maximum intensity projection is done separately on the cortical and cytoplasmic region. To segment the Dvl puncta in the respective region, a 2D Gaussian blur is first performed before a thresholding step. Once the binary image is obtained, the MATLAB function *regionprops* is used to calculate the area and centroid of each puncta. The mean squared displacement with respect to initial position is calculated as 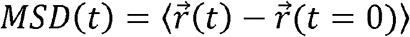, where the average is done over all available puncta. To generate the plot of mean velocity as a function of angle, the instantaneous velocity along the puncta trajectory is first calculated as 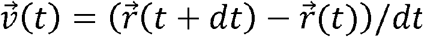, where *dt* is the frame duration. All recorded instantaneous velocities were then binned according to the angle they make with the *x*-axis.

To quantify the fraction of Dvl co-localizing with Lamp1+ endosomes (**Fig. 3F, S4G**), the segmentation of Dvl puncta and Lamp puncta were first performed separately. Briefly, a 2D Gaussian blur was performed before the appropriate thresholding step was applied to obtain a binary image for each channel. The fraction of Dvl on endosome *f*_*Dvl*_ is calculated as *f*_*Dvl*_ = *A*_*Dvl⋂Lamp*_ /*A*_*Dvl*_, where *A*_*Dvl⋂Lamp*_ is the total area of all Dvl puncta that overlap with Lamp puncta, and *A*_*Dvl*_ is the total area of all Dvl puncta. The final reported *f*_*Dvl*_ is the temporal mean averaged over the appropriate cell cycle duration.

Quantification of cortex-associated Dvl signal (**Fig. 4E**) was performed using Fiji [31]. To measure the pixel intensity of Dvl localized to the membrane, a single z-slice encompassing the membrane was selected. A 50-pixel circular region of interest was defined within the vegetal Dvl domain, and a background area was defined in an in-focus region of the oocyte membrane but outside of the Dvl domain. These regions were measured using *RawIntDen* parameter, and the background was subtracted from the Dvl measurement. These background subtracted values were then normalized to the mean of the control oocytes.

## Supplemental Figure Legends

**Supplemental Figure 1.**
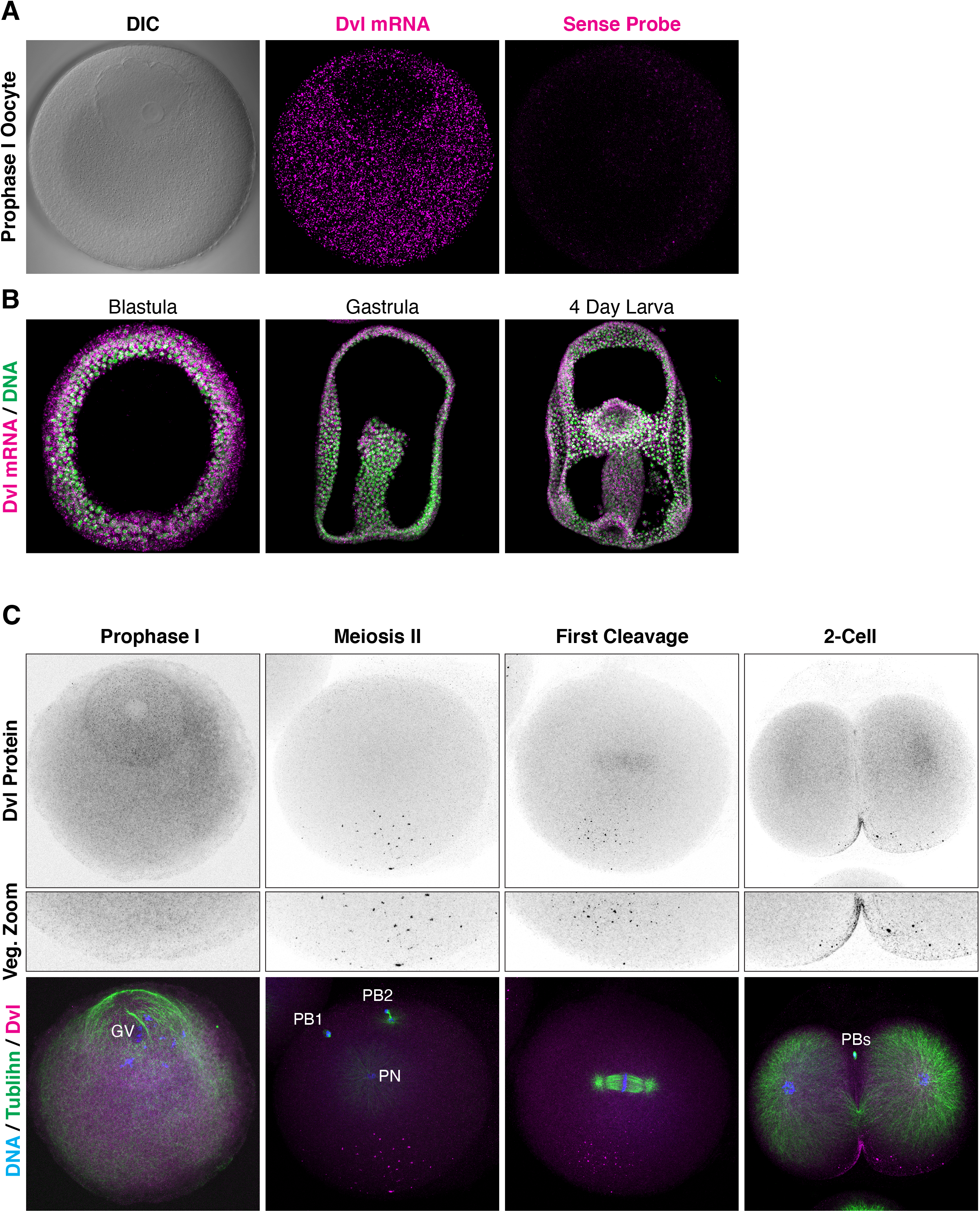
Localization of Dishevelled in meiosis and development, *related to Figure 1*. (A) Fluorescent in situ hybridization for Dvl mRNA in Prophase I arrest of meiosis. Dvl transcripts are detected throughout the oocyte cytoplasm. Sense control probe signal is provided to the right. (B) Localization of Dvl mRNA in embryogenesis. (C) Immunofluorescence for endogenous Dvl protein and microtubules in meiosis and the first embryonic cleavage. Images are scaled individually to better localize Dvl localization.

**Supplemental Figure 2.**
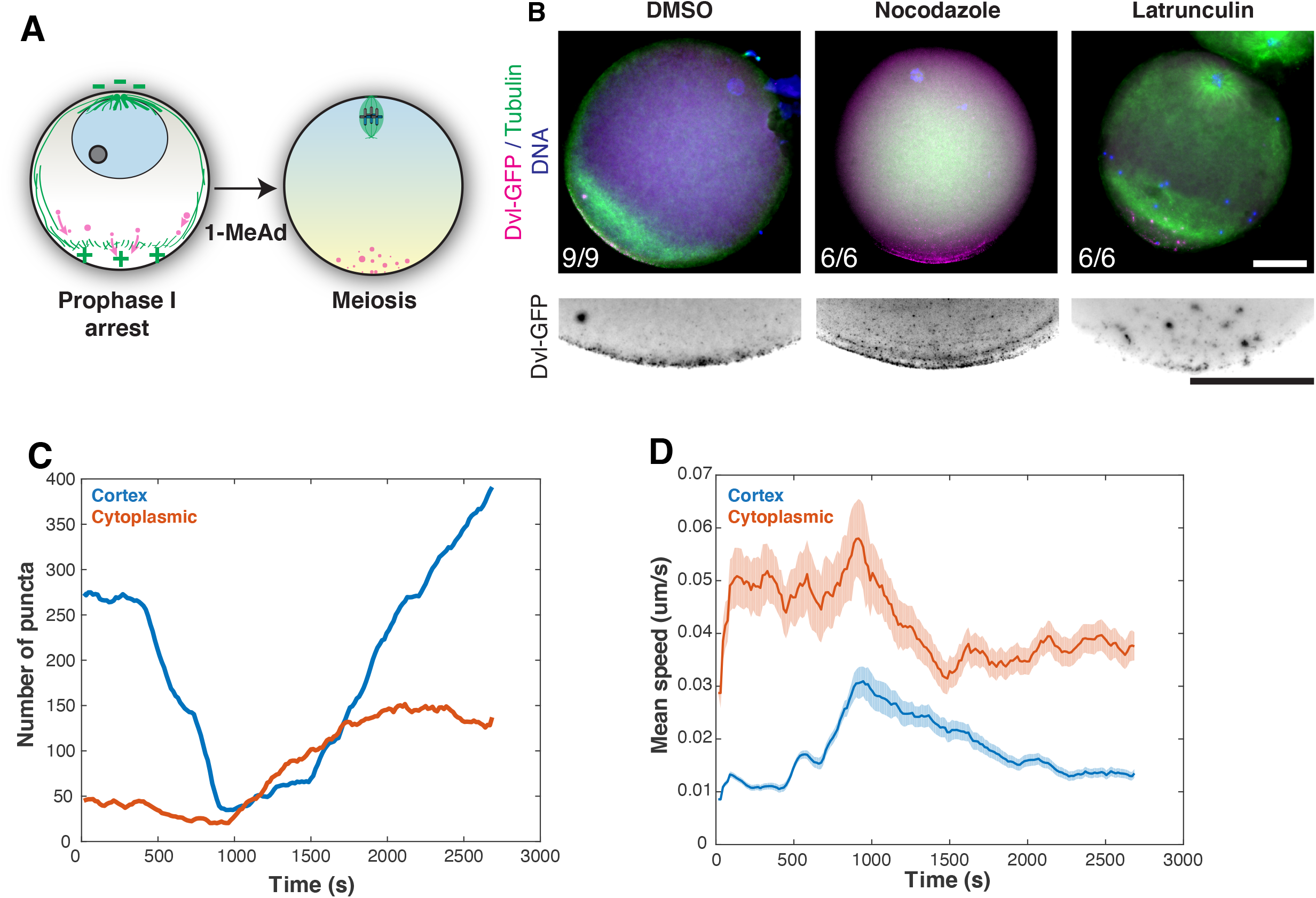
Microtubule independent localization of Dishevelled, *related to Figure 2*. (A) Schematic of a model in which Dvl is actively transported to the vegetal pole along microtubule networks. (B) Fixed oocytes expressing Dvl-GFP and counterstained for microtubules, treated with DMSO control or nocodazole. (C) The number of Dvl assemblies detected over time at the cortex or within the cytoplasm for the oocyte shown in Figure 2A-D. (D) The mean speed of Dvl assemblies at the cortex or in the cytoplasm over time.

**Supplemental Figure 3.**
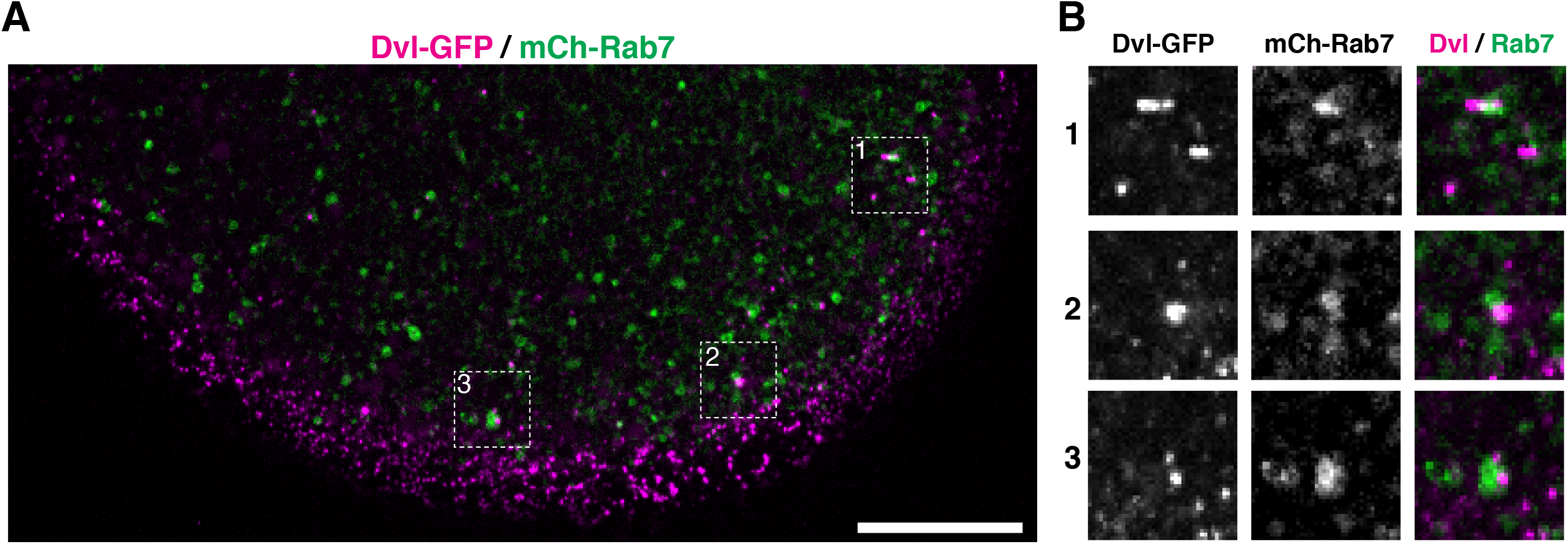
Dishevelled association with Rab7 positive endosomes, *related to Figure 3*. (A) Vegetal pole view of oocyte expressing Dvl-GFP and mCherry-Rab7. Scale bar = 20μ. (B) Zoom views of the insets indicated in (A). Each zoomed image is 8.67μ wide.

**Supplemental Figure 4.**
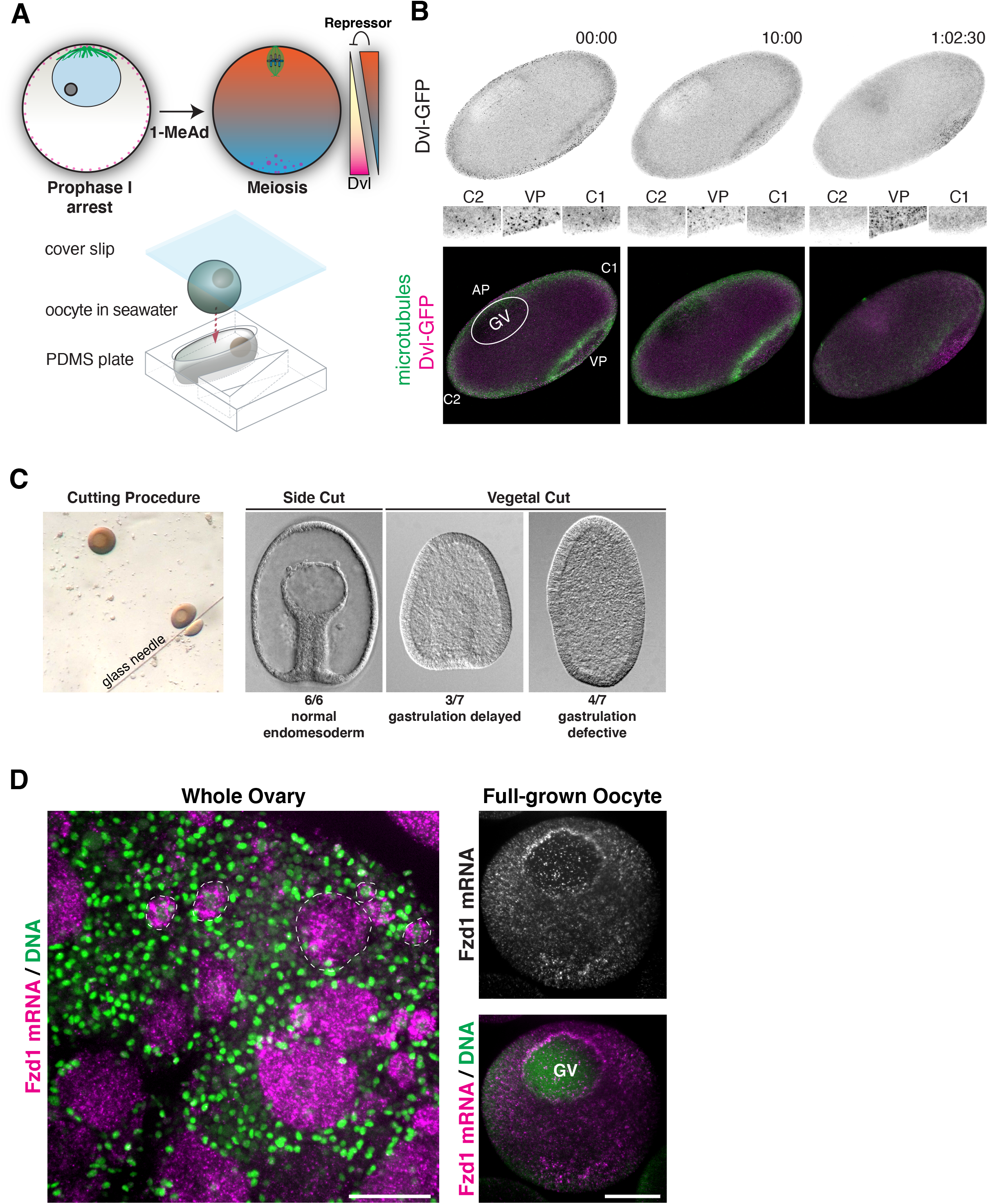
Determinants of Dvl localization, *related to Figure 4*. (A) Schematic of a regulatory gradient model of Dvl localization and PDMS shape mold for modifying the geometry of the oocyte. (B) Time lapse stills of an oocyte in an oval shape mold expressing Dvl-GFP and EMTB-mCherry, representative of 4 individual oocytes. (C) Development of oocytes with either vegetal poles or sides cut as a control at 48 hours post fertilization (control n=6, vegetal pole cut n=7). The cutting procedure by glass needle is shown to the right. (D) Distribution of Fzd1 mRNA in intact ovaries (left) or isolated full-grown oocytes (right). Several example immature oocytes are indicated with white dashed lines. Scale bars = 50μ.

## Supplemental Files

**Supplemental Movie 1. Dvl localization during meiotic resumption**, *related to Figure 1*. Time-lapse video of the oocyte depicted in Fig. 1D,E, with Dvl-GFP (magenta) and the microtubule probe EMTB-mCherry (green).

**Supplemental Movie 2. Dvl dissolution and condensation**, *related to Figure 2*. Time-lapse video of the oocyte vegetal pole at higher magnification, corresponding to the oocyte depicted in Fig. 2A, with Dvl-GFP (magenta) and the microtubule probe EMTB-mCherry (green).

**Supplemental Movie 3. Particle analysis of Dvl assemblies**, *related to Figure 2*. Particles within the cytoplasmic cytoplasmic region (top movie) and cortex-associated region (bottom movie) are tracked separately.

**Supplemental Movie 4. Fusion of cytoplasmic Dvl assemblies**, *related to Figure 3*. Example fusion event between two Dvl assemblies in the vegetal pole cytoplasm.

**Supplemental Movie 5. Association of Dvl assemblies and Lamp1**^**+**^ **endosomes**, *related to Figure 3*. Association of Dvl assemblies (magenta) associated with moving Lamp1^+^ endosomes (green) over time.

## Notes

### Competing Interest Statement

The authors have declared no competing interest.

